# Visual field specializations in mouse dLGN

**DOI:** 10.1101/2025.01.03.631238

**Authors:** Kuwook Cha, Aline Giselle Rangel Olguin, Reza Sharif-Naini, Erik P. Cook, Arjun Krishnaswamy

## Abstract

Neural circuits throughout the visual system process features differently depending on where they appear in the visual field. While such location-specific processing exists in retina and in superior colliculus, the dorsal lateral geniculate nucleus (dLGN) is thought to lack this specialization. Here, we show systematic visual field biases in dLGN’s representation of spatial frequency, orientation, direction, and temporal frequency. Using axon-localized calcium indicators and widefield imaging, we discovered that dLGN boutons show systematic gradients in feature selectivity across the visual cortex (V1), while its retinal inputs lack such gradients for these features. Selective disruption of V1 feedback to dLGN perturbed gradient structure and magnitude. These results suggest that dLGN circuits transform uniformly distributed retinal feature inputs into spatially-biased representations along with cortical feedback. dLGN feature biases would allow a functional stream to detect ethologically salient visual inputs.

## Introduction

Visual scenes are filled with rich detail – some are found anywhere in the visual field, but many are found only in certain locations^1–3^. Think of the way a horizon bisects the visual field or the way textures flow radially when we move. Evolutionary pressures have driven sensory circuits to account for these non-uniformities, and unsurprisingly, visual field specializations are found in many parts of the early visual system^2,3^. In mice, some of these specializations arise in the retina and are retained at later stages^3–12^. For example, visual cortex (V1) inherits retinal specializations for blue and green stimuli in the upper and lower field, respectively^10,13–16^. Others emerge in retinorecipient targets like superior colliculus which enhances certain orientations and motion directions based visual location^17–22^. Similar biases are seen in the V1 of several species^23–29^ and are thought to underlie perceptual biases in humans^28,30^. Thus, location-specific feature processing seems to be a major computation of the early visual system, but an important node, the dorsal lateral geniculate nucleus (dLGN), remains largely unstudied in this regard^31,32^.

The dLGN is the sole link between the retina and V1^33,34^ and its neurons are strongly driven by the axons of retinal ganglion cells (RGCs) – retinal neurons comprising ∼50 different types that encode unique features such as motion or edges^35–38^. Recent advances suggest diverse retinogeniculate connectivity. For instance, slice recordings indicate that some feature-selective RGCs types innervate the same regions of dLGN but never innervate the same dLGN neuron^39^. In vivo recordings show ∼20 functionally distinct dLGN neuron types, each preferring features that mimic that of a single RGC type or is a weighted sum of multiple RGC types^40–45^. This matches single-cell rabies tracings which show dLGN neurons connected to a single RGC type or several RGC types^43,46^. Taken together, these studies suggest that dLGN neurons can enhance or recombine retinal feature signals before delivering this report to V1. Could such dLGN feature computations vary based on visual location?

To address this issue, we delivered axon-localized calcium indicators^47^ (axon-GCaMP8s) to the dLGN and imaged their nerve terminals across the surface and depth of V1. Using widefield imaging, we observed that dLGN boutons exhibit gradients of preferred spatial frequency, orientation, direction and temporal frequency. Two-photon imaging showed that these feature gradients were driven by feature preferences of individual boutons which varied systematically with retinotopic position. But the same experiment detected weak or absent feature gradients across the retinal surface. Finally, genetic ablation of V1-dLGN feedback produced selective deficits in each dLGN feature gradient but did not disrupt them entirely. Taken together, our results show that dLGN circuits weigh inputs from the eye based on retinotopic position to bias its representation of visual features for V1.

## Results

### Spatial organization of visual feature preference in mouse dLGN afferents

While retinal inputs are the main ‘drivers’ of dLGN activity, inputs from V1 also play an important role in ‘modulating’ dLGN signals^33^. To preserve this V1 feedback and measure dLGN feature representation across retinotopic space, we injected dLGN with adeno-associated viruses (AAVs) bearing axon-GCaMP8s^47,48^ (**Fig.1a-b**). Following expression, we implanted mice with cranial windows situated over V1 and widefield imaged GCaMP8s signals from dLGN boutons while mice viewed visual stimuli (**Fig.1c, Supp. Fig. 1a-b**).

**Figure 1.**
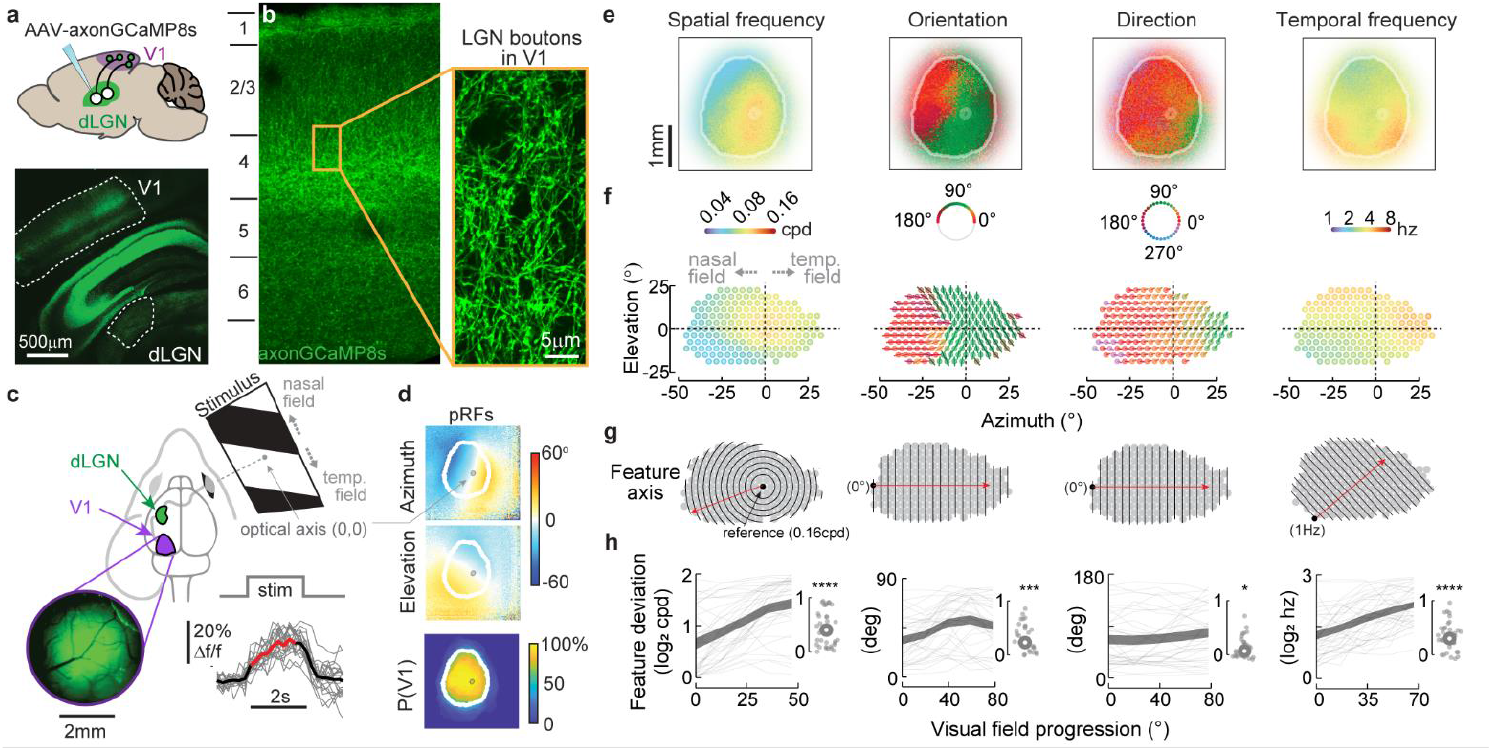
Spatially organized preference for visual features in dLGN inputs to V1. **a**. *Top:* cartoon depicts viral injection of axon-localized GCaMP8s into dLGN to label boutons innervating V1. *Bottom:* confocal image of GCaMP8s expression in a coronal brain slice. dLGN (LGN) and V1 indicated. **b**. Cross section through V1 showing axon-localized GCaMP8s+ dLGN axons. Inset shows a magnified view. **c**. Cartoon depicts a head-fixed mouse viewing stimuli while axon-GCaMP8s fluorescence is collected from a cranial window over V1 (inset). Example single-trial (gray) and average response to a drifting grating are shown. **d**. Standardized azimuth and elevation population receptive field positions (pRFs) and probability map of V1 (P(V1)). White contours delineate the boundary of V1 defined by retinotopy in at least half of the subjects (P(V1)>=0.5). The maps of individual mice were aligned at the region corresponding to the optical axis of the eye (small gray blob). **e**. Average preferred feature maps of dLGN boutons across V1. **f**. Maps from panel e transformed into 2.5° hexagonal bins across visual space. The origin (0,0) is the mouse optical axis. **g**. Axes (red arrow) over which feature preference varies in panel f. At the origin of the axis (black dot), the reference feature from which the feature deviation in h computed is indicated. **h**. Deviation of feature preference with receptive field progression along axes in panel g. Thin gray lines are individuals and thick shaded lines are mean ± standard error (SEM). Inset shows normalized slopes (gray dots) computed from each mouse and the average (gray ring) (n = 36, one-sided t-test; **** for p < 10^−5^; *** for p < 10^−3^; and * for p < 0.05).

We began with a long bar that moved horizontally or vertically along its short axis to map retinotopic organization (**Fig.1c**). Conventional phase and field sign mapping methods^49^ associated dLGN bouton activity with bar position across V1 (**Supp. Fig. 1b**). A small (4-8°) spot flashed at the screen center (**Fig. 1d**, *small gray blob)* let us align and standardize cortical space across subjects and obtain population receptive field positions for each pixel (**Fig. 1d**, *see methods*).

We next presented sets of static gratings, drifting gratings, and full-field luminance-oscillations to measure spatial frequency, orientation, direction, and temporal frequency preferences in the dLGN boutons across V1 (**Supp. Fig. 1a**). Similar stimuli have been used to define feature selective of RGC types^50^. Bouton representations for spatial frequency, orientation, direction and temporal frequency were highly non-uniform across V1 (**Fig. 1e**), suggesting feature preference differs across the visual field. To reveal this organization in visual space, we transformed dLGN bouton feature maps onto the visual field using pixelwise receptive field positions (**Fig. 1f**, *see methods*).

These feature maps showed that dLGN boutons representing the optical axis preferred higher spatial frequencies than those representing more eccentric locations (**Fig.1f**, *spatial frequency*). Boutons responsive to the nasal visual field were biased to horizontal orientations whereas those viewing temporal locations preferred vertical ones with little change in preference across elevation (**Fig.1f**, *orientation*). Posterior motion was preferred at most azimuths except in the temporal visual field where upward motion was preferred (**Fig.1f**, *direction*). Finally, LGN boutons representing higher elevations in the temporal visual field were attuned to higher temporal frequencies as compared to those viewing lower elevations in the nasal field (**Fig.1f**, *temporal frequency*).

To quantify these gradients, we computed the deviation of the feature values of all bins from a reference feature for each map (**Fig. 1g, Supp. Fig. 2a**). This allowed us to express the relative change in feature preference as one progressed across the visual field (**Supp. Fig. 2a-d**). Feature preferences deviated significantly with progression in the receptive field for all four feature maps (**Fig. 1h**); these maps did not result from fisheye distortion (**Supp. Fig. 3**). Thus, we conclude that dLGN feature preferences vary across characteristic gradients across the visual field.

### Feature organization at bouton-level resolution

We next asked how dLGN feature gradients relate to individual bouton activity by two-photon imaging 67 fields from 10 mice across retinotopic positions and depths. Responsive boutons for each stimulus set were selected (73% of 519,603 detected boutons^51^) where only a small set of boutons (16%) were responsive to all 3 stimulus sets, indicating bouton responsiveness is stimulus-specific (**Supp. Fig. 4a)**. Most of the responsive boutons (95%) had their RF within 30° from the population RF location of the field and the remaining 5% were excluded (*see methods*, **Supp. Fig. 4b**).

Example fields representing nasal (Field 1) and central (Field 2) visual locations had similar numbers of GCaMP8s+ boutons but showed strikingly different visual feature preferences (**Fig. 2a-b**). Boutons viewing nasal locations preferred low spatial frequencies, horizontal orientations, posterior motion, and high temporal frequencies (**Fig. 2b-c**, *field 1*). On the other hand, boutons viewing central locations preferred high spatial frequencies, vertical orientations, upward motion, and low temporal frequencies (**Fig. 2b-c**, *field 2*). These results suggested that dLGN boutons viewing similar locations represent similar features, whereas those viewing different locations prefer distinct features.

**Figure 2.**
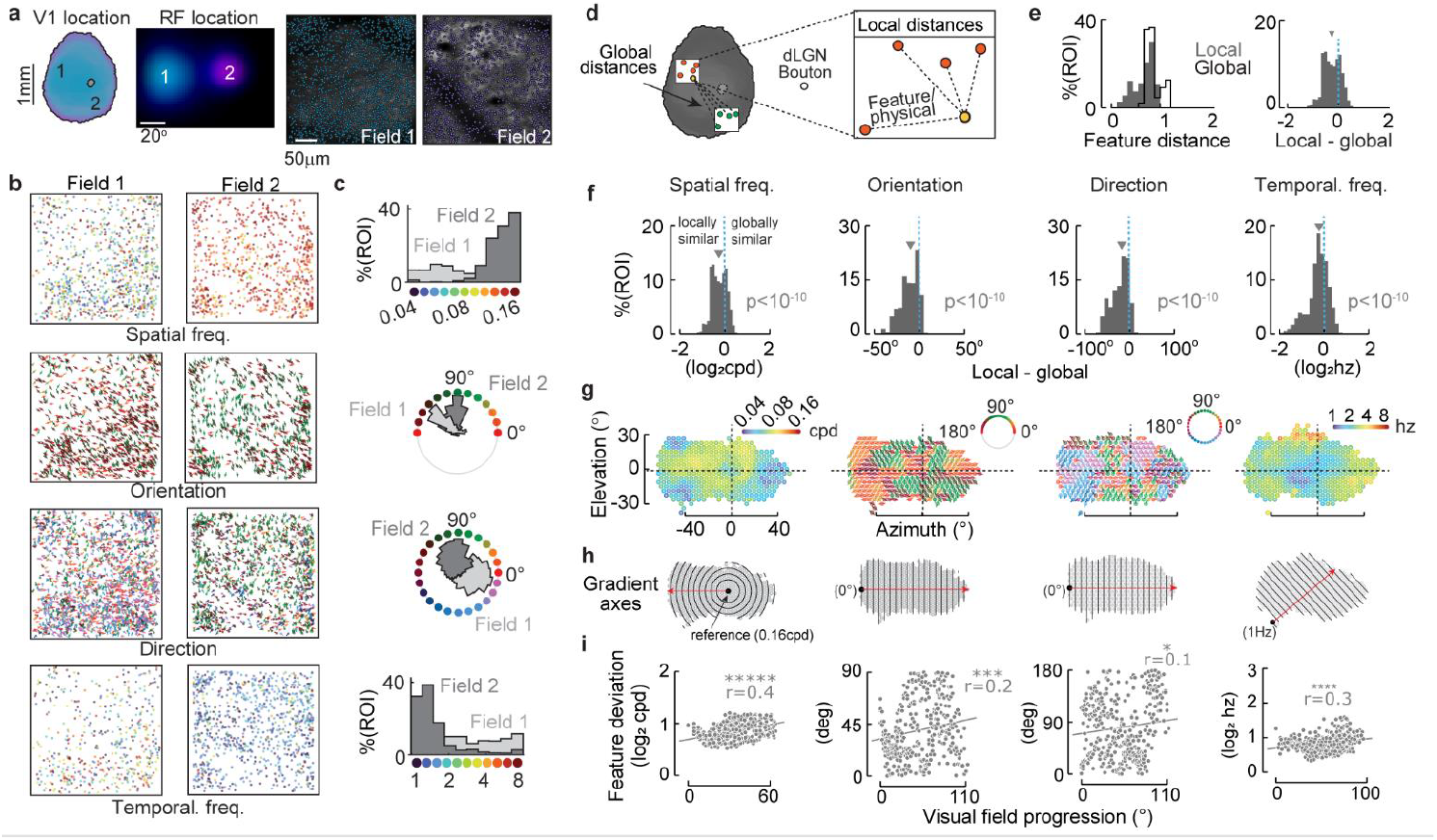
Bouton feature preferences are locally similar and globally organized. **a**. Cartoon of 2 example two-photon fields within V1 (*left*, gray blob corresponds to the optical axis) and their corresponding population receptive fields (*middle*), and average images with responsive bouton ROIs indicated (right). **b-c**. Maps (b) and histograms (c) of preferred spatial frequency, orientation, direction and temporal frequency preference for the fields in panel a. Responsive boutons are stimulus-specific. **d**. Cartoon showing global and local feature distance metrics for a single bouton. The median feature distance between every bouton (yellow dot) and its within-field members (local, orange) was compared to the median distance to boutons in all other fields (global, green dots) **e**. Histogram of local and global feature distances (left) and their difference (right) **f**. Histograms of local-global feature differences for spatial frequency, orientation, direction, and temporal frequency. Distributions are negatively skewed (p < 0.05 in all 4 feature domains, sign-test, n = 67). **g-h**. Bouton feature preferences from all 2p fields binned across the visual field and gradient axes used to measure organization. **i**. Feature deviation versus visual field progression computed from data in panel g using axes in h. Lines are best linear fit. (r = Pearson’s correlation, one-sided test, ***** for p < 10^−10^; **** for p < 10^−5^; *** for p < 10^−3^; and * for p < 0.05).

To confirm this across all visual locations, we quantified similarity in feature preference within the same visual field (*local*) against that across visual fields (*global*). To do this, we computed the median feature distance between a given bouton to its local and global counterparts (**Fig. 2d**). Deducting global from local distance (**Fig. 2e**) showed negatively skewed distributions for all four feature gradients, indicating physically proximate boutons prefer similar features as compared to distant boutons (**Fig. 2f**). This distance-dependent similarity in feature preference continued into a given imaging field such that individual bouton feature preferences were most like their physically proximate neighbors (**Supp. Fig. 4c**). Taken together, these results indicate that feature preference steadily diverges with inter-bouton distance.

Finally, we asked whether individual boutons give rise to feature gradients consistent with those obtained with widefield imaging. To do this, we aggregated and spatially binned bouton data across receptive field positions to create a map covering the entire visual field (**Fig. 2h**). While these feature maps looked more variable than those from widefield imaging, gradient analysis across the axes defined from widefield imaging showed significant trends for all four visual feature representations (**Fig. 2i**). Taken together, we conclude that dLGN feature gradients arise from systematic changes in feature preferences of individual dLGN boutons viewing different parts of visual space.

### Functionally and anatomically defined dLGN bouton subsets drive feature organization

We next asked if some boutons contribute more to gradient structure than others. If true, then feature gradients should be steeper when computed from such bouton subsets. We considered functionally distinct boutons (responsive versus selective) or anatomically distinct ones (across cortical layer).

We started by computing feature maps from either the most responsive boutons for a given stimulus or the most feature-selective ones for a given feature (*see methods*). Gradients for all features were highly significant when computed from the most responsive boutons but were absent when computed from most selective boutons for each feature, except for orientation (**Fig. 3a**). To investigate this further, we computed gradients across overlapping subsets of bouton responsivity and selectivity for each stimulus. Gradients became increasingly pronounced as we moved across increasingly responsive bouton subsets for each feature (**Fig. 3b**) but became increasingly weaker when computed from increasingly selective boutons subsets for each feature. These results indicate that dLGN feature gradients are driven by the most responsive dLGN boutons rather than the most feature-selective ones.

**Figure 3.**
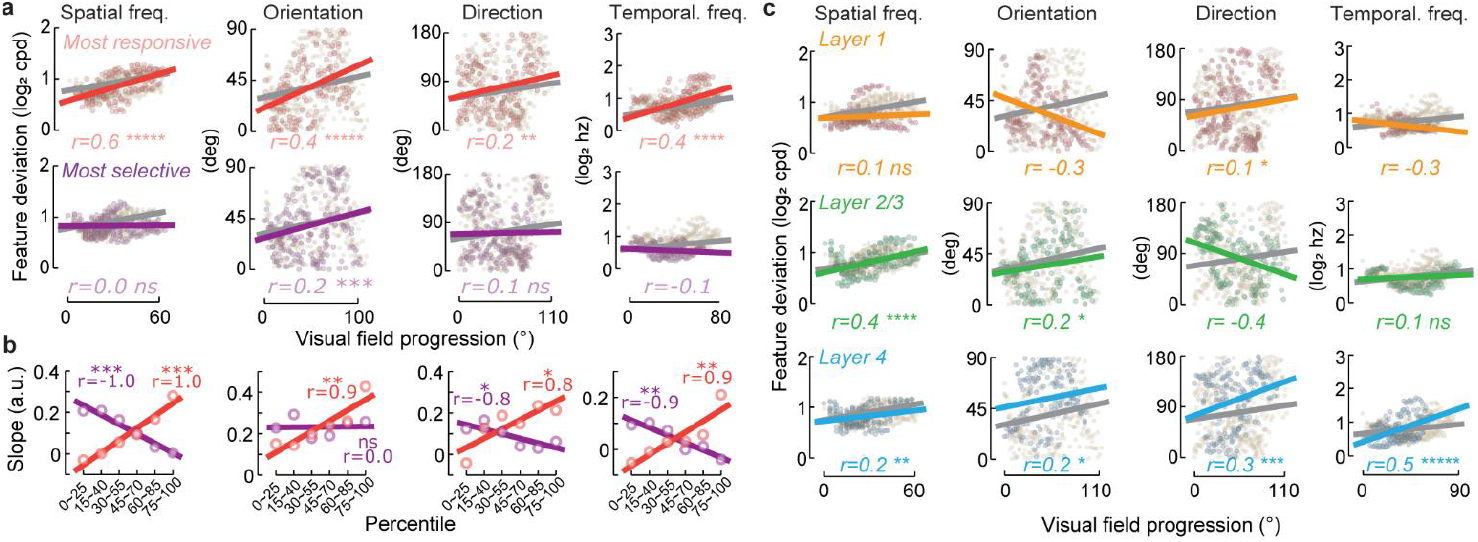
Functionally and anatomically defined dLGN bouton subsets drive feature gradients. **a**. Feature deviation versus visual field progression along the previously indicated axes for most responsive and selective boutons. **b**. Slopes (normalized) from the datasets for the indicated quartiles of responsiveness and selectivity. (*** for p < 10^−3^; ** for p < 10^−2^; * for p < 0.05; and ns for not significant, Pearson correlation test.) **c**. Layer-dependent feature deviation versus visual field progression for most responsive and selective boutons to the indicated visual feature. Lines are linear fits; r, Pearson’s correlation. Significantly positive r-values indicate that preferences change linearly along a feature axis. (***** for p < 10^−10^; **** for p < 10^−5^; *** for p < 10^−3^; ** for p < 10^−2^; * for p < 0.05; and ns for not significant).

Recent work suggests that each cortical layer may be innervated by dLGN neurons carrying a unique mixture of visual information^52–54^. Thus, we next asked if our feature gradients arise from boutons in a specific cortical layer. To test this idea, we computed feature gradients from boutons residing in layer 1 (50-100µm), layer 2/3 (150-250µm), and layer 4 (350-400µm). We saw highly significant gradients for all four features in layer 4 dLGN boutons (**Fig. 3c**, *Layer 4*). Gradients in layer 2/3 boutons appeared only for spatial frequency and orientation (**Fig. 3c**, *Layer 2/3*), and appeared only for direction in layer 1 (**Fig. 3c**, *Layer 1*). Taken together, these results indicate that dLGN feature organization is driven by the most responsive boutons in V1 and is most pronounced for boutons in layer 4.

### Weak feature organization across the retina

Our results so far indicate that dLGN boutons exhibit location-dependent biases in their representation of visual features. Where do these biases emerge? We considered three hypotheses: (1) feature biases first arise from the retina and the dLGN simply relays them to cortex; (2) feature biases require V1 feedback to dLGN; and (3) feature organizations arise in circuitry local to dLGN.

We first tested if dLGN feature organization is inherited from the retina. To do this, we two-photon imaged retinal ganglion cell (RGC) responses to the same stimuli used for our in vivo imaging studies (11,774 identified^55^ RGCs in 9 retinas from 5 mice). Briefly, we explanted retinae from mice in which every RGC expresses GCaMP6f (Vglut2-Cre x Ai62D) and imaged responses from 4-9 fields across each retinal surface (**Fig. 4a-f**). Following imaging, we immunostained retinas with antibodies against Osteopontin which is enriched in dorso-temporal retina and let us align retinas along the cardinal anatomical axes of the eye (**Fig. 4d**). Finally, we used standard methods^56,57^ to map each two-photon imaged field to its immunostained counterpart and transformed these retinal flatmount images back into cups (**Fig. 4c-h**).

**Figure 4.**
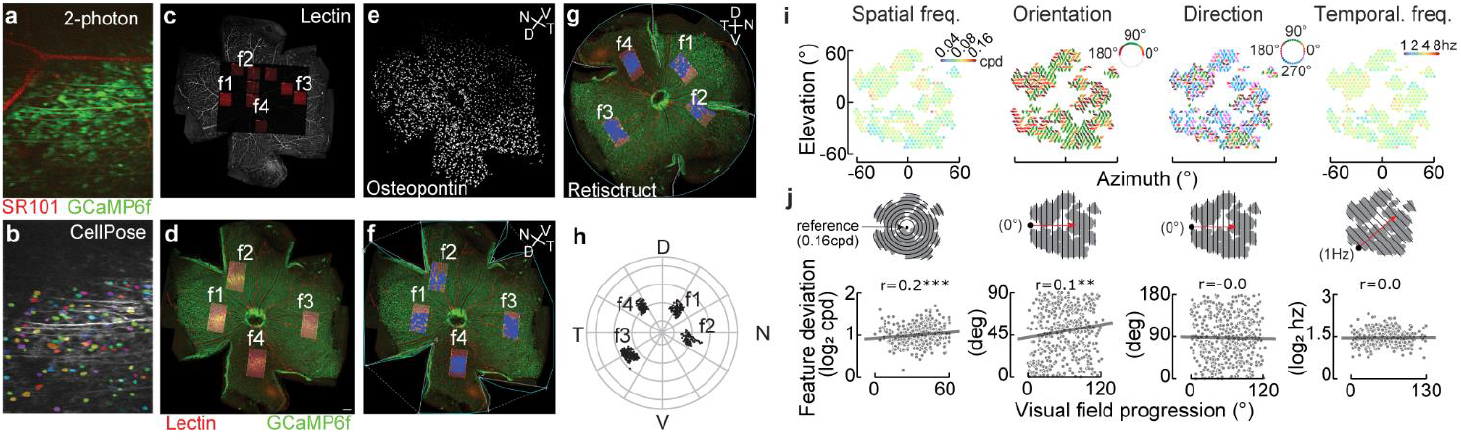
Representation of feature preferences in the Retina. **a**. Example two-photon field of GCaMP6f expressing RGCs counterstained with the blood vessel dye sulforhodamine101 (SR101). **b**. Same field as panel a with RGC ROIs. **c**. Four two-photon imaged fields (f1-f4) aligned to lectin-stained blood vessels. Additional fields were imaged to capture blood vessel patterns for alignment. **d-e**. Two-photon fields mapped onto confocal images of the same retina (d) stained for the dorsotemporal marker, osteopontin (e). N: nasal; T: temporal; D: dorsal; and T: temporal. **f**. Same as panel d but with ROIs (blue dots) for RGCs. **g**. Retistruct transformation of f. **h**. RGC ROIs from g in polar coordinates with the optic disc as the origin. **i**. Binned RGC preferences in visual field space for spatial frequency, orientation, direction, and temporal frequency from experiments like shown in panels a-h. **j**. Feature deviation versus visual field progression along the indicated axes (insets) for the binned RGC data shown in i. Line is best linear fit; r, Pearson’s correlation. Significantly positive r-values indicate that preferences change linearly along the same feature axis as in widefield imaging (*** for p < 10^−3^ and ** for p < 10^−2^).

Next, we spatially binned RGC responses to each visual stimuli as we did for dLGN boutons and mapped each bin to the visual field. Retinal maps of preferred spatial frequency, orientation, direction, and temporal frequency visibly less organized than those of dLGN boutons (**Fig. 4i**). To quantify this, we measured the feature deviation versus visual field progression along the same axes as we did for dLGN boutons (**Fig. 4j**). We saw a small but significant trend for spatial frequency and orientation, suggesting that dLGN may inherit weak biases for these features from the retina. No significant trends along the characteristic axes were seen for direction and temporal frequency. Overall, we conclude that most of the dLGN feature gradients arise after retinal output.

### Contribution of cortical feedback to feature maps

Cortical feedback sharpens and sculpts dLGN visual responses^33,58^. Could these modulatory inputs drive provide the organization of dLGN feature gradients? To address this, we injected V1 layer 6 corticothalamic cell (L6) Cre-line (Ntsr1-Cre^59^) with AAVs carrying Cre-dependent diphtheria toxin receptors (DTR) and labelled dLGN boutons with axon-GCaMP8s (**Fig 5a**). In this way, dLGN feature maps could be imaged before and after chemogenetic ablation of corticothalamic feedback.

**Figure 5.**
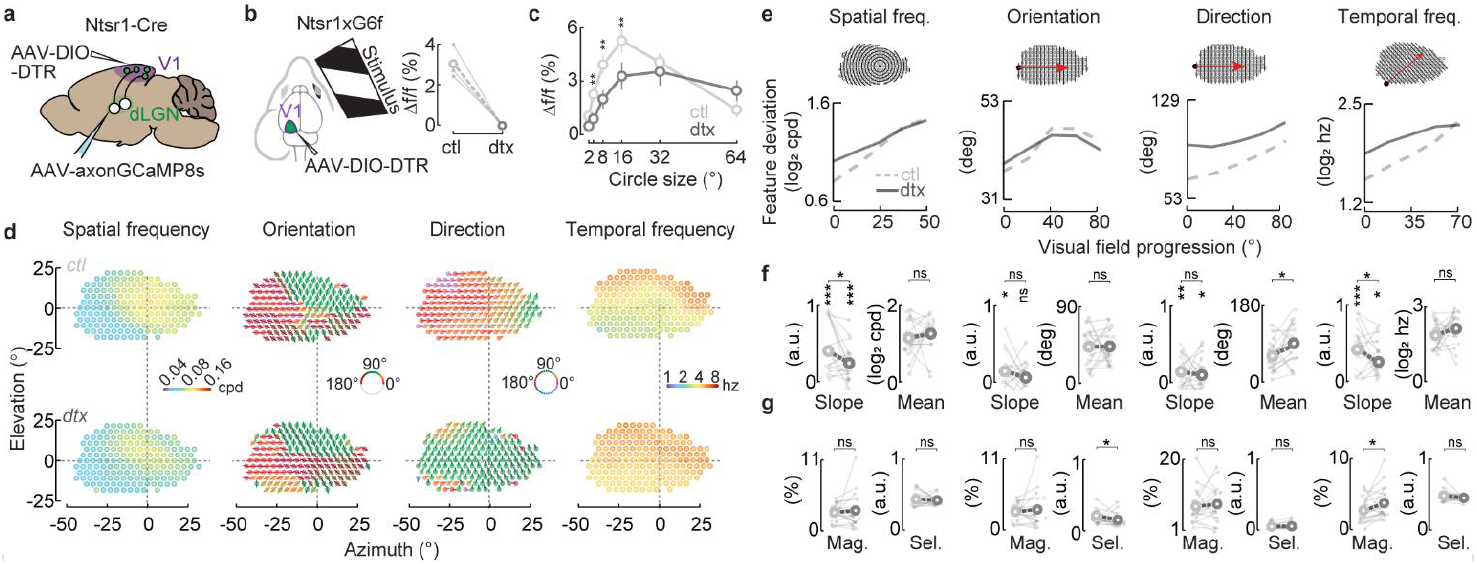
Effects of ablating cortical feedback on dLGN feature maps. **a**. Cartoon of an Ntsr1-Cre mouse injected with AAVs carrying axon-GCaMP8s in dLGN and Cre-dependent diphtheria toxin receptor (DTR) in V1. **b**. Visual responses of GCaMP6f+ layer 6 corticothalamic (L6) neurons widefield imaged from V1 before and after DTX injection. **c**. Epifluorescent responses of GCAMP8s-expressing boutons to an expanding spot before (ctl) and after DTX treatment (dtx). (** for p < 0.01 and ns for not significant, n = 18, paired t-test) **d**. Average bouton feature preferences binned across the visual field before (top) and after (bottom) treatment with DTX. **e**. Feature deviation versus visual field progression along the axis indicated above (inset). Dark gray line (dtx) shows post-ablation average across subjects; light gray, dotted line (ctl), pre-ablation. **f**. Slopes (normalized) and mean values of the linear model fits to individual data (dots) in panel e with the average (rings). (*** for p < 10^−3^; ** for p < 10^−2^; * for p < 0.05; and ns for not significant, t-test, n = 18). **g**. Overall response magnitude (Mag) and feature selectivity (Sel) computed over the entire V1 for data in d. Dots, individual data; rings, condition average. (* for p < 0.05, n = 18, paired t-test).

The effects of chemogenetic ablation were confirmed in two ways. First, we saw a complete loss of visual responses in L6 neurons themselves 5 days after DTX injection in a separate group of mice that expressed GCaMP6f and DTR in these cells (**Fig. 5b**). Second, dLGN bouton responses to a disc with varying size showed significantly altered size tuning after DTX ablation of L6 neurons, consistent with prior reports of feedback modulation of dLGN responses^58^ (**Fig. 5c**).

We next asked how dLGN feature gradients were affected by loss of L6 feedback. Prior to DTX treatment, feature gradients were consistent with our earlier findings (**Fig. 5d**, *top*). Post-ablation maps generally resembled the control data (**Fig. 5d**, *bottom*), but each feature representation showed selective alterations following treatment with DTX (**Fig. 5d-g**). While the dLGN spatial frequency gradient remained following DTX treatment, the steepness of this gradient was significantly reduced (**Fig. 5e-f**, *Slope*). The overall response magnitude (**Fig 5g**, *Mag*) and feature selectivity (**Fig 5g**, *Sel*) remained the same, indicating the gradient reduction was not caused by disrupted or absent responses to this stimulus. dLGN orientation gradients were no longer significant after DTX treatment despite some remaining organization indicated in the average trend (**Fig. 5e-f**, *slope*). Consistent with this, we also saw an overall reduction in orientation selectivity after DTX (**Fig. 5g**). Direction gradients remained significant and did not differ from the control condition (**Fig. 5e-f**, *Slope*) but there was a global rotation in the preferred direction relative to their pre-injection counterparts (**Fig. 5f**, *Mean*). There was however no associated change in overall response magnitude or selectivity for direction (**Fig. 5g**). Temporal frequency gradients were also retained but were significantly shallower than controls (**Fig. 5e-f**, *Slope*). Overall temporal frequency selectivity did not alter, but there was an increase in response magnitude which might be related to temporal precision modulated by cortical feedback^60^ (**Fig. 5g**). Taken together, these results indicate that L6 feedback sharpens each dLGN feature gradient but does not create them entirely.

## Discussion

Here, we investigated whether mouse dLGN weighs visual features according to location in the visual field. By widefield imaging axon-GCaMP8s+ dLGN boutons across V1, we observed characteristic gradients of spatial frequency, orientation, direction, and temporal frequency preference across the visual field. Two-photon imaging showed that these gradients arose from systematic changes in highly localized feature preferences of dLGN boutons viewing different visual locations. Retinal imaging showed weak or absent biases, and loss of corticogeniculate feedback produced selective deficits in feature gradients but did not ablate them entirely. Our results demonstrate visual field specializations in dLGN, and suggest they arise from dLGN circuits that integrate V1 feedback.

### More than just a relay

Growing evidence indicates that dLGN neurons connect to RGC types in specific ways to either enhance or recombine retinal feature signals^39–46^. Since dLGN boutons arise from neurons residing in different sectors of dLGN and do not receive direct synapses in V1, we reason that our feature gradients reflect systematic changes in the feature preferences of dLGN cells.

Prior work showed feature-preferring dLGN neurons^40–45^ and indicated their preferences arise from synapses formed with specific RGC types^39,43,46^. By inferring patterns of somatic activity from boutons, our results indicate that such wiring specificity can vary depending on where a neuron resides in the dLGN. Importantly, our results predict that neurons residing in the posterior dorsomedial dLGN (nasal visual field) should connect specifically with horizontal orientation-selective RGCs (OS-RGCs)^61^; and those in the anterior ventrolateral region (temporal field) should connect with vertical OS-RGCs^35,61,62^.

Innervation biases have been seen with horizontal and vertical OS-RGC projections to the superior colliculus (SC) whose orientation maps strongly mirror those we have observed in dLGN^18^. Given that most RGCs innervating dLGN collateralize to innervate SC^63^ and given that mechanisms for retinotopic targeting are shared between both structures^64–66^, it is possible that this pattern of OS-RGC axon innervation is also shared. Imaging the OS-RGC boutons across dLGN through cannula-implants or retrograde tracing from posterior dorsomedial and anterior ventrolateral dLGN into retina could shed more light on this issue.

### Cortical feedback enhances dLGN feature maps

V1 can strongly modulate response properties of dLGN neurons^33^. Studies in several species show various effects including surround modulation^58,67,68^ and temporal precision^60,69^, alteration in feature preference^70–72^, and attentional modulation in the presence of distractors ^73^. Although corticothalamic projections modulate local circuits in the thalamus, L6 neurons might also directly influence geniculate boutons through translaminar inhibition^59^.

While the loss of cortical feedback impacted every dLGN feature map, the effects were specific to each representation (**Fig. 5**). Systematic changes in spatial and temporal frequency preference remained following ablation of layer 6 feedback, but the strength of this gradient was substantially weaker. Azimuthal gradients in orientation preference were lost following ablation and were accompanied by generally weaker orientation selectivity. Gradients for motion-direction remained but appeared to rotate overall preference from posterior to vertical motion. Why were these effects feature specific?

One idea comes from reports in NHPs showing that corticogeniculate feedback is feature- and stream-specific^67,74^. The differing effects we observed in our ablation studies could reflect the loss of a few kinds of L6 feedback cell types. Recent studies suggesting distinct L6 corticothalamic types in mice^75–77^ strongly support this notion. Such differing L6 types could enable feature-specific enhancements. For example, spatial frequency tuning might be modulated through surround suppression^58^ while the altered orientation maps and the global rotation effects in motion direction could involve cortical neurons preferring specific features^70–72^. Finally, changes in temporal frequency maps can be attributed to corticothalamic control of temporal precision^60,69^.

### Functional specialization and visual ecology

While most visual features are thought to have an equal likelihood of occurring across a natural scene, some features depart from this ‘translation invariance’. Blue skies fill the upper visual field of many terrestrial vertebrates, whereas their lower field contains earthier tones^7^. The biases we have shown in our results along with prior work on visual field specializations could be viewed as evolutionary adaptations to visuo-ecological niches. While prior studies speculated that such specializations belong to more ancient parts of the early visual pathway (e.g., retina and SC), our work suggests that specializations are present at the level of dLGN. What purpose could they serve?

One clue could be that dLGN feature maps are driven by the most responsive boutons as opposed to the most selective ones. This might reflect the presence of two streams of dLGN input to V1 – a strongly driven stream that is biased by visual location, and a more selective stream with weaker biases. The notion of parallel channels has been suggested by recent studies that different dLGN types project to different layers in V1^43,54,78^. Perhaps a ‘biased’ stream enables fast perceptual judgements that capture ethologically salient features with other subcortical pathways like SC, whereas the ‘unbiased’ stream is used for versatile scene analysis. Accordingly, it has been proposed that different visual tasks, such as detection versus discrimination, involve extracting visual information from distinct neural populations^79–81^. Training mice to detect or discriminate features, such as oriented lines, in different locations across the visual field while providing distractors that either match or mismatch dLGN feature maps could shed more light on the behavioral significance of the specializations we have described here.

## Contributions

KC and AK, conceived of this study. KC, AK, and EC aided with instrumentation, analysis, experiments, writing, and making figures. AGRO performed retinal experiments. RS provided important experimental advice and comments on the manuscript.

## Acknowledgements

We thank J. Forestell, X. Ma, S. Sharif, J. Lehnert, N. Brake, and F. Jalondoni for helpful comments on the manuscript.

## Methods

### Animals

Male and female wildtype C57BL/6, Ntsr1-Cre (RRID:IMSR_JAX:030264) and VGluT2-Cre mice (RRID:IMSR_JAX:030264) were used. For wide-field experiments, 18 wildtype animals and 18 Ntsr1-Cre animals were used. Eight of these 36 mice and 2 additional mice were used in 2-photon imaging experiments. Rosa26-CAG-GCaMP6f (RRID: IMSR_JAX:028865) mice crossed with VGluT2-Cre mice for experiments with RGCs. All surgical and experimental procedures were in accordance with the rules and regulations established by the Canadian Council on Animal Care, and protocols were approved by the Animal Care Committee at McGill University.

### Virus and diphtheria toxin injections

Mice were anesthetized during virus injections at 2% isoflurane (4% for induction). A Neuros syringe (Hamilton) was used to deliver AAV-hSyn-axon-jGCaMP8s-P2A-mRuby3 (Neurophotonics, Laval) to dLGN. To prevent spill-over into the visual cortex, we approached dLGN from the top of somatosensory area (-2.1mm lateral and 0.8mm posterior to the bregma) at a 30-degree angle, targeting 2.1mm lateral and 2.6mm posterior to the bregma and 3.1 mm below the skull. Injection was performed slowly (200-250nl for 3-4 minutes) using a microinjector (World Precision Instruments) and the needle was kept in position for >5 minutes before it was retracted.

Using the same setup, AAV2/retro-CAG-flex-DTR (Neurophotonics) was delivered to the left V1 in Ntsr1-Cre mice. For wide expression over the V1, injection was performed at 3 sites which are 0.5∼1mm from the center of the V1 (2.5mm lateral and 0.1 anterior to the lambda) 600µm below the dura. A volume of 150nl was delivered to each site. Usually, these injections were done at cranial window implantation. Diphtheria toxin (Sigma D0564) was administered intraperitoneally (100 ng/g body weight) to mice expressing DTR after their control imaging session.

### Headplate and cranial window implantations

Mice were induced with 4% isoflurane and analgesics (carprofen at 5cc/g and topical lidocaine) given before surgery. Anesthesia was maintained using 2% isoflurane. After surgery, the animal was placed back in its home cage and provided with carprofen for the following 2 days.

Headplates were implanted after dLGN injections over the exposed skull around the left V1 using dental cement. For this, a circle was marked for the placement of cranial window and dental cement was applied to the surrounding area. A headplate with a circular hole was placed on top of the cement and then additional dental cement was applied. After recovery, the animal underwent cranial window surgery. A 4 mm-diameter craniotomy was made over the left V1 using a dental drill. After removing the skull flap, a round cover glass was gently placed in the craniotomy and sealed with cyanoacrylate glue skull while kept in place with a small rod attached to a micromanipulator.

### Histology

Euthanasia of mice by isoflurane overdose and perfusion with phosphate buffered saline (PBS) was followed by brain extraction. The brains were postfixed in 4% (w/v) paraformaldehyde (PFA) overnight. A tissue slice (Compresstome) was used to collect 150- or 200-um thick coronal sections of the brain. For immunostaining, the brain sections were incubated in blocking buffer (10% normal donkey serum, 0.4% Triton X-100 in PBS) for 1-2 hours, followed by incubation in primary antibodies at 4°C for 7 days, and secondary antibodies at 4°C overnight. Sections were imaged under a Zeiss confocal microscope where 405nm wavelength was used for excitation.

Two-photon imaged retinas were dissected and fixed for 45 min in chilled 4% (w/v) PFA, 20 min in methanol and 20 min in acetone. For immunostaining whole-mounts were incubated with blocking solution (4% normal donkey serum, 0.4% Triton-X-100 in PBS) for 2 hours, followed by incubation with primary antibodies overnight at 4°C. Tissue was then washed with PBS and incubated in secondary antibodies for 2 hours at room temperature. Images of stained tissue were acquired on a Zeiss LSM-710 inverted confocal microscope at a resolution of 512 × 512 pixels with a 0.3 μm step size.

Antibodies used were as follows: chicken anti-GFP (1:1000, Abcam, Cambridge, UK; RRID:AB_300798) and goat anti-osteopontin (1:1000, R&D Systems, Minneapolis, MN; RRID:AB_2194992). Secondary antibodies were conjugated to Alexa Fluor 488 (Cedarlane, Ontario, CA; RRID:AB_2340375) or Cy3 (MilliporeSigma; RRID:AB_92570). Isolectin (Fisher Scientific, Waltham, MA; RRID:SCR_014365) was incubated along with the secondary antibodies.

### Visual stimulation

Visual stimuli were presented on an LED screen (34.4 cm × 19.4 cm) placed perpendicularly to the optical axis of the right eye of the mouse. The center of the screen was located 10 cm away from the optical axis and the left end of the screen was aligned to the midline of the subject. This resulted in a coverage of 120 degrees in azimuth and 80 degrees in elevation. The red color channel of the monitor was disabled, and all stimuli were presented on a linearized monochromic scale out of a uniform background at the midpoint luminance (grey).

Visual stimuli were implemented with Psychtoolbox and controlled by in-house software run on MATLAB. For retinotopic mapping, a white or checkerboard bar swept across the screen either horizontally or vertically at the speed of 10 degrees/second. The thickness of the bar was 10 degrees. Each direction was repeated 40 times for wide-field imaging and 10 times for 2-photon imaging. For size tuning, a white or black disc was presented at the center of the screen with the diameter randomly chosen each trial from a set of 2, 4, 8, 16, 32 and 64 degrees. Each of 120 presentations was 2-second long and planked by grey background screen (1 second before and 2 seconds after presentation).

For feature tuning, static gratings, drifting gratings and luminance-oscillations were presented for 2 seconds each trial subtending the full screen (**Supp. Fig. 1a**). In static grating experiments, spatial frequency was randomly chosen out of 0.04, 0.08 and 0.16 cycles/degree, and orientation was drawn from 0, 45, 90 and 135 degrees. The spatial phase was also randomly chosen from 0, 90, 180 and 270 degrees. Typically, 240 trials were given for widefield imaging and 120 for 2-photon imaging. In drifting grating experiments, one of 8 directions (0, 45, 90, 135, 180, 225, 270, and 315 degrees) was drawn each trial. The spatial frequency was set at 0.05 cycles/degree and the grating drifted at the speed of 60 degrees/second (3 Hz). Typically, 160 trials were given in widefield imaging and 80 for 2-photon imaging. In luminance-oscillation experiments, spatially uniform luminance changed sinusoidally in time for 2 seconds each trial at one of 4 frequencies (1, 2, 4, and 8 hertz). The temporal phase was either 0 or 180 degrees, randomly drawn each trial. Typically, 160 trials were delivered in widefield imaging and 80 for 2-photon imaging.

We did retinal stimulation as described previously^57,82–85^. Briefly, a DLP light crafter (Texas Instruments, Dallas, TX) was used to project dichromatic (405nm, 520 nm) visual stimuli through a custom lens assembly that steered stimulus patterns into the back of a 16× objective. All visual stimuli were displayed with a background intensity set to 1 × 104 R*/rod/s. The projector LED and the scan retrace of the two-photon microscope was synchronized using custom electronics^86^. The 3 feature stimulus sets (static gratings, drifting gratings and full-field luminance oscillations) were presented with a spatial conversion at 30um per degree. Each stimulus was presented in a fixed order for 1 second twice. A 1-second-long gray background period was given between presentations.

### In vivo wide-field and two-photon imaging

For widefield imaging, we used a custom-built epi-fluorescent microscope to capture the visual cortex (V1) through a cranial window at a 10hz sampling rate using custom code written in LabVIEW (National Instruments). Mice were lightly sedated (0.5-1% isoflurane) on a semi-cylindrical platform with its headplate affixed to a metal holder. Acquired images covered a 4 mm×4mm field of view at a 180×180 resolution.

For two-photon imaging, cranial-window implanted mice were placed under a custom-built galvo-galvo-resonant two photon microscope and lightly sedated. A 16x NA .8 water immersion objective (Olympus) was lowered atop the cranial window to focus into V1. The fluorophores were excited at 920 nm using a Mai-Tai (Newport) equipped with dispersion compensation and laser power at the sample plane was typically 5-15 mW. A 280×280 um^2^ field of view was scanned at 45Hz as a series of 512×512 images. Imaging depth was measured from the pia which was identified by illuminating the field at 1024nm. Imaging field locations were determined by real-time observing active boutons responding to a bar sweeping across the visual field. Once an imaging field location is decided, it was attempted to image three depths of layer 1 (50-100um), layer 2/3 (150-250um), and layer 4 (300-350) along the same column. Collecting data from all 3 layers was possible in sixteen field locations (48 fields), which were included in the layer-specific feature analysis. An additional 19 fields were included in all the other analyses.

### Two-photon imaging of retinal ganglion cells

Two-photon imaging was performed as previously described^57,82,83,85,87–89^. Briefly, mice were dark adapted for at least 2 hours, euthanized, and then retinas rapidly dissected under infrared illumination into oxygenated Ames solution (95% O2, 5% CO2; MilliporeSigma, A1420). Next, retinas were mounted onto a filter paper (MilliporeSigma, HABG01300) with the RGC layer facing up, placed in a recording chamber, mounted on the stage of a custom-built two-photon microscope, and perfused with oxygenated Ames solution warmed to 32–34°C. Responses of GCaMP6f+ RGCs to visual stimuli delivered through the objective were imaged at 920 nm (MPB Femto fixed 920nm laser) and collected at an imaging rate of 45 Hz. Each image plane (483.5×483.5um) of the movie contained GCaMP fluorescence, SR101 fluorescence, stage coordinates, and visual stimulus synchronization pulses to permit offline analysis. A few microliters of sulphorhodamine 101 (SR101, 2 mg/mL, MilliporeSigma, S7635) was added to the recording chamber to label blood vessels and a map of the main blood vessels emanating from the optic disk acquired for post hoc image registration. Data acquisition was performed with ScanImage synced to the visual stim using a photodiode sync pulse and handshake pulse using custom made code in MATLAB. Following recording, retinas were fixed, immunostained and imaged as described above.

### Time-series data processing for in vivo cortical imaging data

The following time-series processing was applied to each pixel of widefield imaging data and each ROI of cortical 2-photon imaging data. Suite2p software^51^ was used to define ROIs (519,603 in total) and extract their time-series data from 2-photon images. Time-lapsed image intensity values of each trial were first resampled at 5hz and were converted to proportion of change from a baseline, ΔF/F = (F - F_0_)/F_0_. The baseline, F_0,_ was computed from the average values of a 1-second-long grey background period that preceded stimulus onset each trial. These values underwent a Gaussian smoothing over 5 consecutive trials to increase reliability of the estimation and alleviate the effect of transient changes. In the case of bar stimuli for retinotopy, the time series data were concatenated across trials for Fourier analysis. For the other stimuli, the response for each trial was obtained by averaging ΔF/F over the period of stimulus presentation (for 2 seconds after stimulus onset).

### Data processing of retinal imaging data

ROIs were defined using CellPose^55^, and MATLAB was used to extract time series data for each ROI. The signals were smoothed to remove noise with moving mean of 10 frames and expressed as ΔF/F, where the mean of the peristimulus baseline was used to calculate the difference. The response for each stimulus presentation was obtained by taking the maximum value of the calcium signal during the 1-second of stimulus presentation. Responsiveness of ROIs were evaluated for each of the three stimulus sets as in cortical image data analysis (See above) except the response reliability was computed as the maximum response divided by the standard deviation of baseline activity. Only ROIs whose response is above 3 standard deviations were included in the feature analysis.

Retinas were immunostained after two-photon calcium imaging and image registration was performed as previously described^57^. Briefly, the blood vessel image of each recorded field was stitched into a single image using custom MATLAB scripts, while a confocal image was acquired for the same field after staining. Following scaling, rotation, and translation the two images were overlayed to match sulforhodamine101 and lectin blood vessel patterns. Next, fine blood vessel morphology was used as inputs for landmark correspondence in Fiji to register each two-photon field into a whole-mount retina confocal image. The whole-mounted retina was orientated in the D-V-T-N axis using the density of alpha RGCs as a reference (higher density in the D-T region) which were labelled with an osteopontin staining. The orientation of the stimuli presented during two-photon imaging was also orientated using alpha RGC density as reference.

Following registration, each ROI obtained was mapped into the whole-mount retina space using the linear transformation matrix obtained from Fiji’s landmark correspondence. These RIOs were projected from a flattened retina into a reconstructed standard spherical retina space using the Retistruct package for azimuthal equilateral projections. All ROIs from 9 retinas were merged in single projection, where left retinas were mirrored into the right retina space. Polar coordinates given by Retistruct were then transformed to cartesian coordinates of visual field positions centered at the optical axis to match the coordinate system of the in-vivo data.

### Widefield retinotopy mapping and image alignment

Each pixel of widefield image data from retinotopy experiments underwent fast Fourier transform to estimate the phase of the periodic responses to bar positions sweeping across the screen horizontally or vertically (**Supp. Fig. 2b**). The azimuth and elevation phase representations in the image space were used to compute vector field gradients to find the boundary of V1 in each mouse.

A cortical region that is responsive to the center of the screen/the optical axis of the eye was defined for alignment of image data across individual mice in the following way. Single-trial responses to a small circular stimulus (4 or 8 degrees in diameter) presented at the center of the screen were converted to t-values and the t-map was smoothed (Gaussian filter with a sigma value of 4 pixels). A blob of spatially clustered pixels with the highest 0.2 % t-values (∼65 pixels) was defined as the cortical representation of the optical axis, whose center was used as a reference point to be aligned to across mice. Image translation was performed to align the images at the reference point. We refer to the aligned image space as standard image space in which all individual mouse datasets reside as a result of the alignment.

After the alignment, the primary visual cortex was demarcated by selecting the pixels that were indicated as the primary visual cortex in at least 18 of the 36 mice (P(V1)>=0.5). This region we refer to as standard V1 is used for all 36 individual datasets. Further data analysis was done only in the pixels of this region.

In the cortical ablation experiments, within-subject image registration was performed using MATLAB image processing toolbox to align datasets acquired before and after ablation. Mean images from the first run of the retinotopy imaging were used as a reference image.

### Bouton ROI selection

After Suite2p selected tentative bouton ROIs, responsiveness of the Suite2p ROIs was evaluated for each of the three stimulus sets (static gratings, drifting gratings, and full-field luminance oscillations). Single-trial responses of the preferred stimulus condition, which was determined by the largest average response of all conditions, were t-tested and ROIs that passed a criterion of p < 0.01 were selected for further analysis. The selection was performed for each stimulus set so that different sets of ROIs were chosen for analysis (**Supp. Fig. 4a**). The t-scores of the preferred response of these ROIs were also used in the analysis for responsive quantiles.

The second criterion was the deviation of receptive field position from the median receptive field position of the boutons in the same imaging field. We examined the overall distribution of the deviation and set the cutoff value at 95% which corresponded to 30.1 degrees (**Supp. Fig. 4b**). Therefore, ROIs whose receptive field position was away from the population median of the field farther than 30.1 degrees were discarded.

### Estimation of receptive field position

Receptive field position was estimated in the same manner for each pixel of widefield data and ROIs of in vivo two-photon data. The position of the bars sweeping across the screen either horizontally or vertically was resampled at the same rate as time series (5 Hz) and the average across trials was approximated on a grid of every 2.5 degrees. The fluorescence time series data were also averaged across trials over one condition (e.g., horizontal sweeps), which yielded a ‘position tuning’ function. The position tuning function was interpolated at every 0.01 degree and the peak position was assigned as receptive field position. Azimuth and elevation were estimated separately.

After estimation of pixelwise (i.e., population) receptive field position in widefield data of each mouse, the median across the individuals were computed in the standard image space. There was a systematic shift in these estimates indicated by the receptive field position of the screen center region in widefield data, which was (6.3, 6.5). This shift, presumably caused by the delay in fluorescence responses, was corrected by subtracting the offset values. In the case of widefield data, the resultant azimuth and elevation values, we refer to as standard azimuth and elevation, were assigned back to individual data.

### Computation of preferred feature and feature selectivity

The following analysis was applied to each pixel of widefield data and each bouton ROI in 2-photon data, similarly to a previous study^54^. Preferred features were estimated using a vector sum formula, 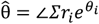, where r_i_ and θ_i_ are the average response and the feature value for stimulus condition i. For orientation, θ_i_ was 2 times of the stimulus orientation angle and 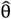 was multiplied by 0.5 as preferred orientation. In the case of spatial frequency and orientation, since the 2 feature domains were combined in a static grating stimulus, the best stimulus parameter of the other domain was determined first. For example, in estimation of preferred orientation, the best spatial frequency for a pixel or an ROI was first found and the responses to orientations were chosen among the responses to that spatial frequency. The same principle was applied to estimating preferred spatial frequency.

For spatial frequency and temporal frequency, the original feature values were converted to angular values on a semi-circle. In other words, spatial frequencies 0.04, 0.08 and 0.16 cpd were log-scaled and evenly placed in the semi-circular space, resulting in 0, π/2 and π. In the same way, temporal frequencies 1, 2, 4, and 8 Hz resulted in 0, π/3, 2π/3 and π, respectively. Vector averaging on a semi-circle leads to compression of the resulting values towards the centroid. We overcame this by adjusting r_i_ so that the minimum is set to 0, which allowed the preferred feature value to be distributed across the full range of the feature space. After the vector averaging, 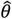 on the semi-circle was converted to the native frequency values by inverse mapping. Feature selectivity was computed by utilizing the circular operation as well: 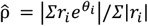.

In averaging preferred feature values either across animals or within a spatial bin, spatial frequency and temporal frequency were log-transformed, averaged and transformed back to the native frequency scale; and orientation and direction were averaged using the vector sum method described above.

### Spatial binning

Spatial binning was applied to create feature maps in the visual field space and feature gradient analysis. Pixelwise data from widefield imaging were spatially binned based on receptive field positions into a hexagonal grid. The radius of the bin was set as 2.5 degrees. The same grid was applied to in vivo 2-photon ROIs. In the case of dLGN bouton ROIs, only the bins with at least 30 ROIs were used for further analysis. In the case of 2-photon data from the retina, only the bins with at least 10 ROIs were included.

### Feature gradient models

Systematic feature preference changes were captured as gradients of the changes along characteristic axes. Of note, this approach is analogous to previous studies that used consistent positive changes across positional segments to quantify similar phenomena; but quantifying them as a single slope measure is a more stringent and succinct method.

Feature gradient axes were determined by observation of the visual field maps of preferred features and did not involve numerical optimization. The gradient axis is expressed as follows: (1) for spatial frequency, 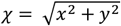 (2) for orientation, *χ* = *x*, (3) for direction: *χ* = *x*, and (4) for temporal frequency, 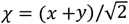, where x is azimuth and y is elevation. The origin of the x-y coordinate system corresponded to the optical axis of the eye.

The gradient model is generally described as *g*(*χ*) = *β*_0_ + *χβ*_1_ + *ε*, where *g*(*χ*) is feature deviation from a reference value at *χ, β*_0_ and *β*_1_ are the intercept and the slope of a linear fit, and *ε* is an error term. The feature deviation was defined for each domain as follows(**Supp. Fig. 2a**): (1) for spatial frequency, *g*(*χ*) = | *log*_2_θ(*χ*) − *log*_2_0.16| where θ(*χ*) is spatial frequency at *χ*; (2) for orientation, *g*(*χ*) = |*angdiff*(θ(*χ*), 0)| where *angdiff* is angular difference; (3) for direction, *g*(*χ*) = |*angdiff*(θ(*χ*), 0)|; and (4) for temporal frequency, *g*(*χ*) = | *log*_2_θ(*χ*) − *log*_2_1|. The receptive field positions of widefield data were transformed to *χ* and the feature values were transformed to *g*, so a linear function could be fit to estimate *β*s for each mouse, using the least-squares method (MATLAB polyfit function). Slope estimates *β*_1_ were obtained in each mouse (**Supp. Fig. 2b-d**) and underwent a t-test. They were normalized for visualization in Fig. 1g and Fig. 5g so that the maximum possible slope value would be 1. For 2-photon imaging data, Pearson’s correlation was computed to test the linear relationship between *χ* and *g*.

## Supplemental Figures

**Figure S1.**
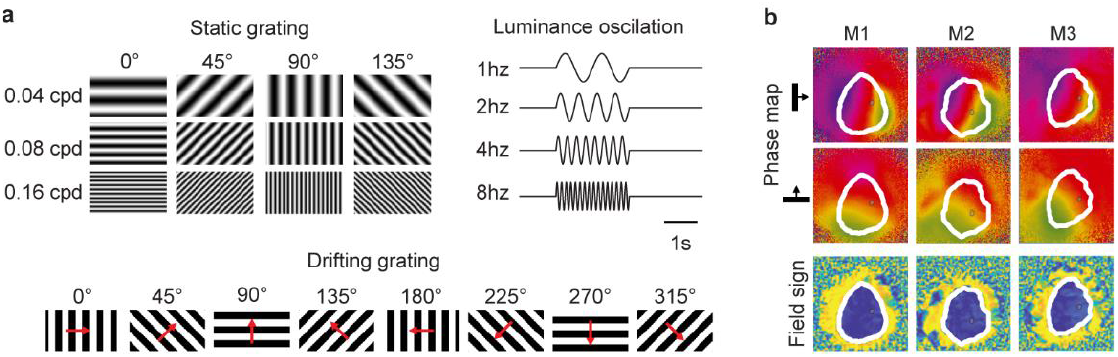
Feature stimulus sets and retinotopic mapping. **a**. The static grating set (top right) included 12 conditions of 3 spatial frequencies (0.04, 0.08 and 0.16 cpd) x 4 orientations (0, 45, 90 and 135°) conditions. Spatial phase was randomly drawn out of 4 (0, 90, 180, and 270°) each presentation. The drifting grating set (bottom) included conditions of 8 directions (0, 45, 90, 135, 180, 225, 270, and 315°). The grating drifted at 60°/s and the initial phase was randomly drawn from 0, 90, 180, and 270°. The luminance oscillation set included 4 conditions of temporal frequencies (1, 2, 4 and 8 Hz) and the initial phase was randomly drawn from 0 and 180°. **b**. Retinotopic phase maps and field sign maps of 3 example mice. White blobs represent V1 defined by field signs, and small gray blobs represent regions responsive to the center of the screen. All image data across subjects were aligned at the center of the blob.

**Figure S2.**
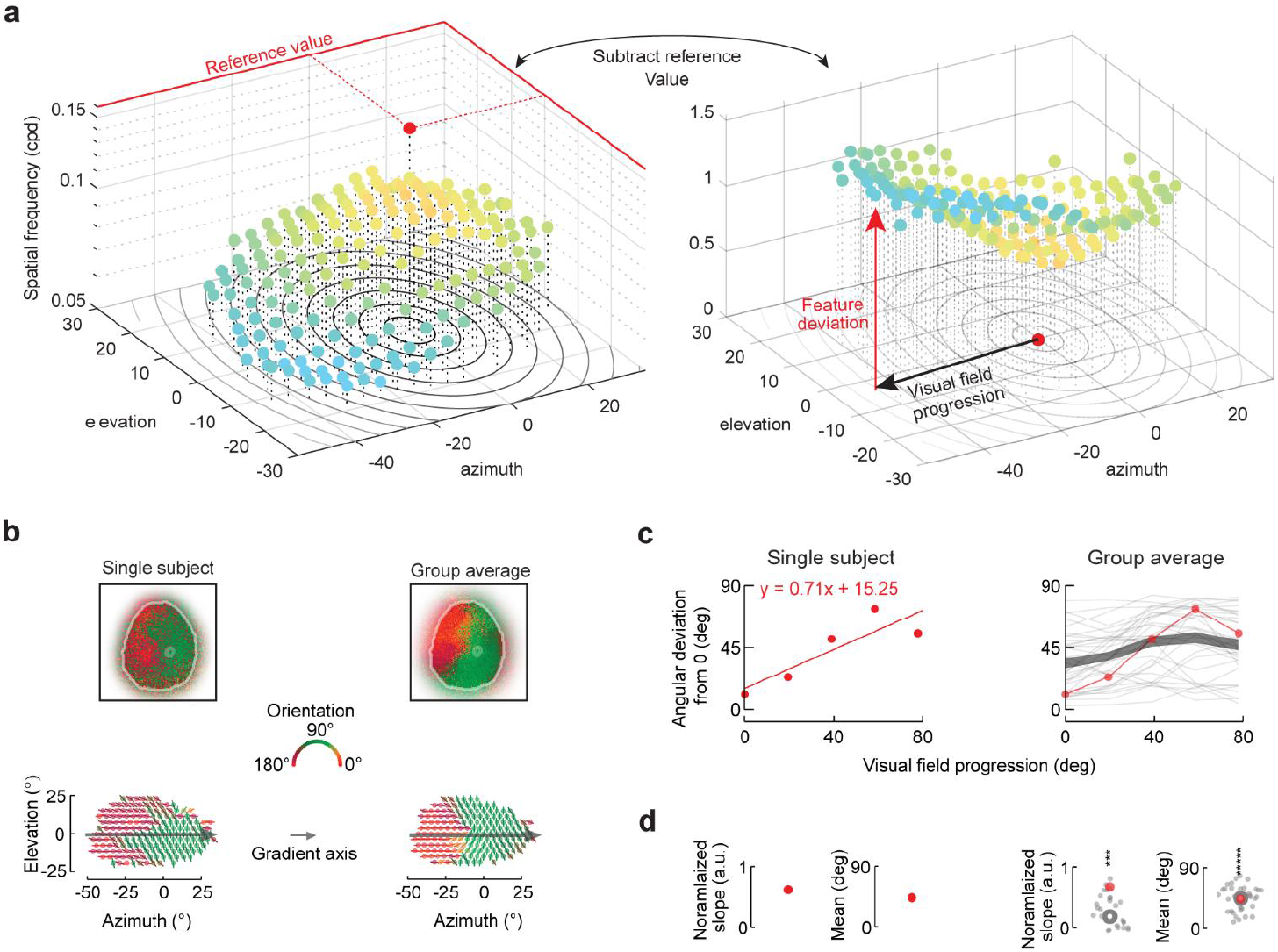
Gradient analysis. **a**. 3D scatter plot of spatial frequency preference binned across the visual field. A reference value is deducted from each bin to transform the map into a gradient plot allowing measure of visual field progression and feature deviation. **b**. Example (left) and average (right) orientation maps on visual cortex (top) and visual field (bottom). Arrow indicates the naso-temporal gradient axis for orientation. **c**. Left: single subject data binned across the gradient axis (red dots) and fit with a line. Right: the same binned data also shown with other subjects (red and gray thin lines) and group average (Thick line mean ± SEM). **d**. The slope and the mean estimates of the linear model in a single subject (left) and plotted alongside all animals (right). Average slope and mean estimates indicated.

**Figure S3.**
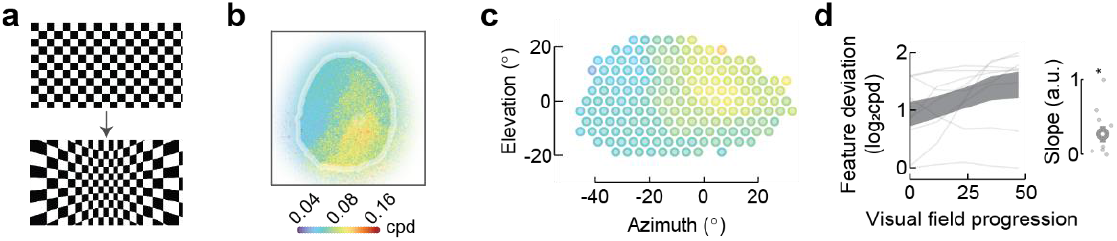
Spatial frequency gradient after fisheye distortion correction. **a**. Checkerboard image before (top) and after (bottom) fisheye compensation. A subset of mice (n = 9) underwent a separate experiment using static grating stimuli with this image transform. **b**. Average map of spatial frequency preference over cortex. **c**. Visual field representation of b. **d**. Gradient along eccentricity and the normalized slope (inset). Shaded area, mean ± stand error. Thin lines, individual mouse data. In the inset, small dots are individual data; and open circle, mean. * indicates p < 0.05 (Wilcoxon signed rank test, one-sided, n = 9).

**Figure S4.**
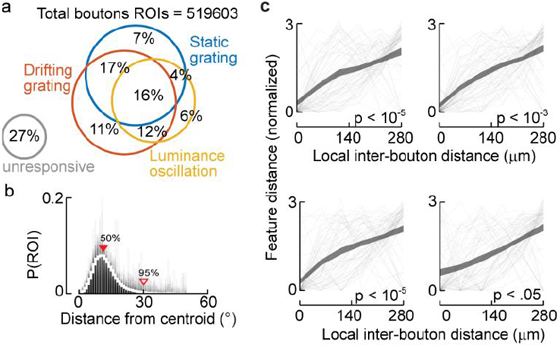
Bouton selection and similarity within and across two-photon fields. **a**. Proportions of responsive boutons out of 519,603 regions of interest (ROI) identified by Suite2p. Responsiveness was tested with t-tests for the response of each ROI to their preferred stimulus within each stimulus set. Each of the 3 ROI sets was used for the respective feature analysis. **b**. Scatter of receptive field (RF) measured by deviation (distance) from the population RF center of the field. Only responsive ROIs were included (N = 379,311). The population RF center was computed as the median location of ROIs within the field. Histograms of each field are overlaid in gray and the white contour represents the average distribution across all fields. Triangles indicate 50- and 95-percentiles. Most boutons (95%) fall within about 30 degrees in their RF from the centroid, and the ROIs with their RF outside this distance were excluded in the feature analysis. **c**. Monotonic increase in feature distance as a function of inter-bouton distance within the field. Data from 67 fields are plotted as thin lines overlaid by a thick line for mean ± SEM. Slopes of linear fit to individual data were significantly above zero (p < 0.05 in all 4 feature domains, t-test, n = 67).

